# sCentInDB: A database of essential oil chemical profiles of Indian medicinal plants

**DOI:** 10.1101/2025.02.24.639916

**Authors:** Shanmuga Priya Baskaran, Geetha Ranganathan, Ajaya Kumar Sahoo, Kishan Kumar, Jayalakshmi Amaresan, Kundhanathan Ramesh, R.P. Vivek-Ananth, Areejit Samal

## Abstract

Essential oils are complex mixtures of volatile compounds produced by aromatic plants. Due to their odor and therapeutic properties, essential oils are used in personal care, food flavoring and pharmaceutical industries. As a high-value and low-volume organic product, optimizing plant yield and modifying composition by leveraging knowledge on chemical profiles of essential oils can lead to enhanced bioproducts. To this end, we present sCentInDB, a manually curated **D**ata**B**ase of E**s**sential oil **C**hemical profil**e**s of Medici**n**al plan**t**s of **In**dia, which has been built based on information in published literature. sCentInDB is a FAIR-compliant database which compiles information on 554 Indian medicinal plants at the plant part level, encompassing 2170 essential oil profiles, 3420 chemicals, 471 plant-part-therapeutic use associations, 120 plant-part-odor associations, and 218 plant-part-color associations. For the documented essential oils, sCentInDB also compiles extensive metadata including sample location, isolation and analysis methods. Subsequently, an extensive analysis of the chemical space of essential oils documented in sCentInDB was performed. By constructing a chemical similarity network, the chemical space was found to be structurally diverse with terpenoids distributed across the network. Moreover, a comparison of the scaffold diversity in sCentInDB was performed against three other aroma libraries using cyclic system retrieval curves. Altogether, sCentInDB will serve as a valuable resource for researchers working on plant volatiles and employing genetic engineering to enhance oil yield and composition. Further, sCentInDB will aid in establishment of quality standards for essential oils and provide vital insights for therapeutic and perfumery applications. sCentInDB is accessible at https://cb.imsc.res.in/scentindb/.

## Introduction

Essential oils are complex mixtures of volatile compounds produced by plants, and are characterized by strong odor and therapeutic properties.^1–3^ These volatile substances are secreted and accumulated through specialized anatomical structures.^2^ The term “essential” reflects the fact that it encapsulates the fragrance essence of the plant.^4^ Essential oils are commonly obtained from various plant parts via distillation or a mechanical process.^5^ These oils play a crucial role as internal messengers, in plant defense, plant-plant interactions, and attracting pollinators.^2,4,6–8^ Notably, essential oils have been reported to exhibit antimicrobial, anti-inflammatory, and anticancer properties, and therapeutic effect against cardiovascular diseases and other health conditions.^9–15^ Due to their distinct odor and therapeutic properties, essential oils have found applications in perfumery, personal care, pharmaceuticals, and food and cosmetics industries.^16–20^ In particular, these oils have been traditionally used in aromatherapy to promote health and holistic well-being.^1,21^ India, with its rich biodiversity and numerous endemic plant species, has a long-standing tradition of utilizing medicinal plants in various traditional systems of medicine.^22,23^ While significant research has been conducted to characterize essential oils from Indian medicinal plants^24,25^, much of this knowledge remains scattered across the literature, underscoring the need for its systematic compilation and utilization.

Typically, essential oils consist of chemical components with low molecular weight and volatility at ambient temperature.^1,26,27^ These oils contain specific classes of chemicals, including terpenoids, shikimates, polyketides and lipids, produced through major pathways of plant secondary metabolism.^2,28,29^ Variations in these chemical components are influenced by factors such as plant developmental stage, temperature, water availability, and climatic conditions, which contribute to the unique composition, yield and aroma of the oil.^5,21,30–33^ The chemical constituents of the oil are generally categorized into major and minor components, with the minor components typically comprising less than 5% of the total composition.^34^ Though the major chemical components of the oil primarily determine its biological properties, minor components can also play a crucial role by exerting antagonistic, additive, or synergistic effects, thereby influencing the oil’s overall bioactivity.^14,35^ Notably, these minor components can also greatly contribute to the odor of the oil.^36^

Chemical profiling of an essential oil is crucial for ensuring the quality standards^37^, identifying the odorous chemicals responsible for oil fragrance^38,39^, optimizing the yield and composition of the oil through biotechnology application^40^, and determining the suitability of the oil for medicinal use or as food flavoring.^2,41,42^ Essential oils are high-value, low-volume bioproducts due to low plant yields, and hence changing the yield and composition of essential oils is important ^43–45^. Additionally, modifying their composition helps reduce antinutritional factors ^46^ or undesirable components, thereby improving the overall quality and usability of the oils. While chemical synthesis and microbial cultures can produce essential oil constituents, they are limited by high costs^47^, structural complexity of phytochemicals^48^, and gaps in understanding cellular processes^49^, making biotechnology-based methods a promising alternative. To this end, tissue culture techniques help to genetically improve and obtain fully grown, mature plant with desirable traits.^40,50,51^ Additionally, modified plant cell cultures are often utilized to produce secondary metabolites of interest, such as terpenoids.^51^ These approaches require detailed knowledge of the chemical composition and biological activities of the oil.

To date, a few databases have been developed to catalog the chemical profiles of essential oils. However, these resources are not region-specific or do not focus exclusively on medicinal plants, leading to limited coverage of medicinal plant species. Among the notable databases, EssOilDB^4^ and AromaDB^52^ provide valuable information on essential oils. EssOilDB^4^ was the first publicly available resource containing information on essential oils from plants worldwide, curated from published research articles. AromaDB^52^ specifically captures the aroma molecules in essential oils of medicinal and aromatic plants. Apart from these freely accessible databases, a few proprietary resources such as EOUdb (https://essentialoils.org/plans) and AromaWeb (https://www.aromaweb.com/) have been developed to provide information on essential oils and aromatherapy. However, no dedicated effort has been made to develop an essential oil database for Indian medicinal plants, in particular, one that focuses on oil data derived from plant samples collected within India.

In this contribution, we compiled from published literature, the chemical components of essential oils from Indian medicinal plants, with a particular focus on plants whose part samples were also sourced within India. Additionally, we gathered data on their therapeutic uses, odor, and color from published literature. To assist chemists in utilizing this data, we annotated the chemical compounds with relevant information such as physicochemical properties, drug-likeness characteristics, ADME properties, and molecular descriptors. The compiled resource represents the most comprehensive database to date in terms of plant coverage and detailed chemical information on essential oil-bearing medicinal plants from India, providing a valuable knowledgebase for scientific research. The compiled database is accessible via the associated website: https://cb.imsc.res.in/scentindb/.

## Methods

### Data collection

To compile the chemical profiles pertaining to essential oils of Indian medicinal plants, we manually curated information from 778 published research articles. In order to search the published literature for essential oil profiles, the list of Indian medicinal plants was compiled from the IMPPAT database^53,54^, information provided by the Ministry of AYUSH, Government of India, and books on the subject. This resulted in a non-redundant list of around 7500 Indian medicinal plants. Thereafter, published articles providing chemical profiles of essential oils of Indian medicinal plants, were searched from various sources including PubMed (https://pubmed.ncbi.nlm.nih.gov/) and Google Scholar (https://scholar.google.com/). This search for articles was conducted until March 2022. From the identified literature sources, information on plant part from which the essential oil was extracted, location from which the sample was collected, isolation method, analysis method, chemical constituents, retention index, retention time, and percentage of the individual chemicals in an essential oil, were collected. During this compilation, we included only essential oil data from Indian medicinal plants, where the location of the plant sample collection was reported to be within India. In few instances, the chemical composition of the essential oil in terms of individual chemicals summed up to more than 100% (as reported in the published source), and such inconsistent information were omitted in this compilation. Furthermore, information on the therapeutic uses, color, odor, and general uses of an essential oil were compiled from the corresponding published source along with additional literature sources and books. In sum, through this extensive manual effort, we curated chemical profiles of essential oils from Indian medicinal plants, focusing exclusively on plant samples collected within India (Figure 1). Figure 1 summarizes the workflow of this study, from literature mining to compilation, curation and harmonization of the data contained in sCentInDB.

**Figure 1:**
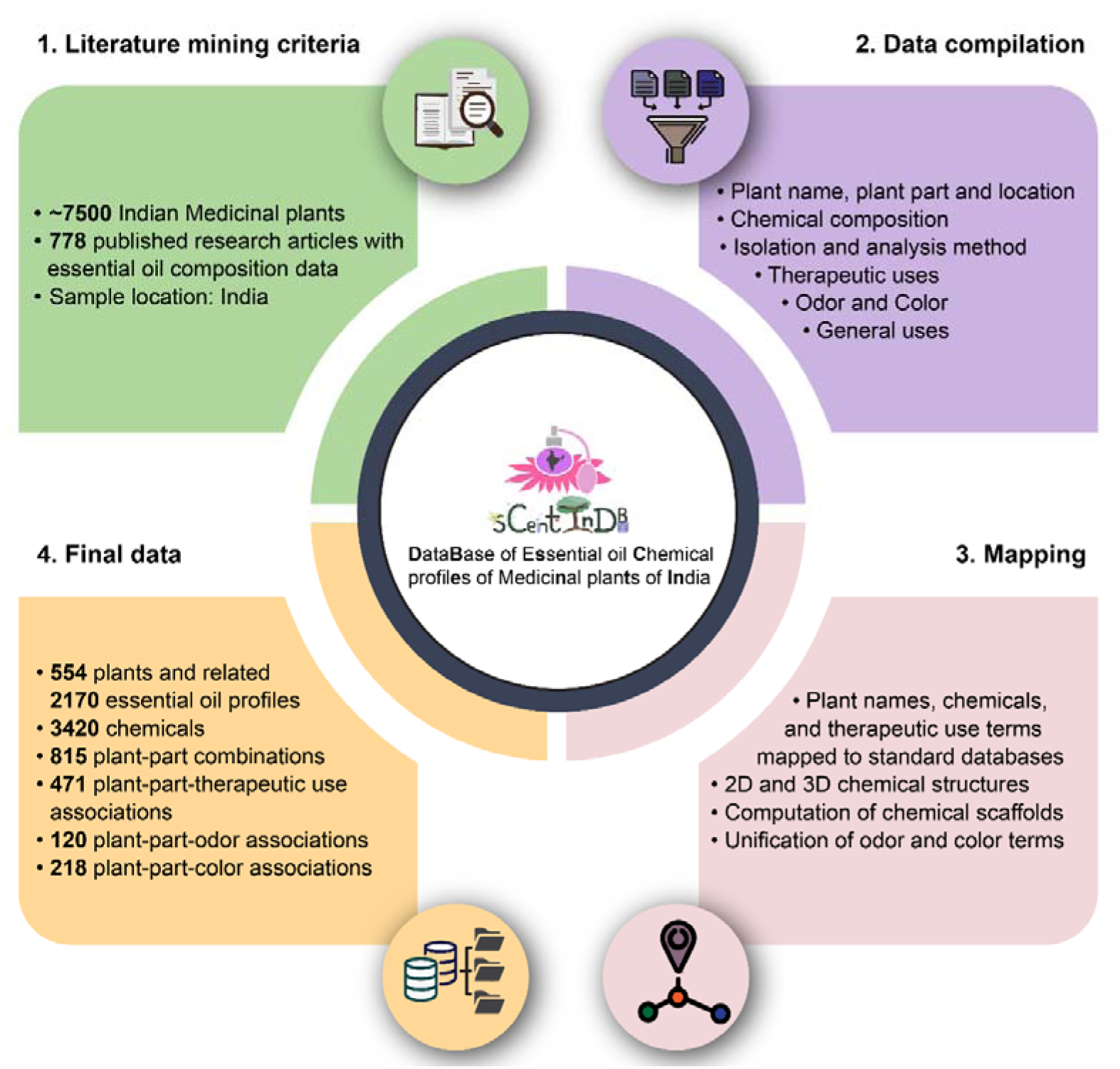
Workflow summarizing the entire process of sCentInDB database construction involving Literature mining, Data compilation, Mapping and annotation, and Final dataset.

### Plant annotation

We mapped the reported names of medicinal plants with essential oil profiles to their corresponding accepted scientific names using three resources namely, The World Flora Online (WFO) (https://www.worldfloraonline.org/), Plants of the World Online (https://powo.science.kew.org/), and Tropicos (https://tropicos.org/). This resulted in a non-redundant list of 554 medicinal plants with essential oil profiles in this compiled dataset. Furthermore, common names, taxonomic information and conservation status or extinction risk of these medicinal plants were compiled from Flowers of India (https://www.flowersofindia.net/), WFO (https://www.worldfloraonline.org/) and IUCN Red List (https://www.iucnredlist.org/en), respectively. Moreover, information on the use of medicinal plants in various traditional Indian systems of medicine, such as Ayurveda and Siddha, was compiled from the IMPPAT database. Additionally, for these medicinal plants with reported essential oil profiles, we provided links to other resources namely, WFO (https://www.worldfloraonline.org/), Tropicos (https://tropicos.org/), Medicinal Plant Name Services (https://mpns.science.kew.org/), Plants of the World Online (https://powo.science.kew.org/), International Plant Names Index (https://www.ipni.org/) and Gardeners’ World (https://www.gardenersworld.com/).

### Annotation of essential oil chemicals

We obtained a non-redundant list of 3420 chemicals across essential oil profiles compiled in sCentInDB. The two-dimensional (2D) structure of the essential oil chemicals were obtained from PubChem^55^ (https://pubchem.ncbi.nlm.nih.gov/), ChemIDplus (https://pubchem.ncbi.nlm.nih.gov/source/ChemIDplus) and ChemSpider (https://www.chemspider.com/) databases. For chemicals that could not be mapped to information in these chemical databases, we manually generated their 2D structures using the Chemicalize Marvin JS (https://chemicalize.com) software, based on the structural representations provided in the corresponding published research articles or books. Moreover, the 2D structure information of chemicals was stored in various formats such as SMILES, InChI, InChIKey, SDF, MOL and MOL2, which were generated using Open Babel.^56^ For the chemicals, the images of their 2D structure were generated using RDKit (https://www.rdkit.org/). Further, the three-dimensional (3D) structures of the chemicals were retrieved from PubChem. If the 3D structure of a chemical was not available in PubChem, the 3D structure was generated from its 2D structure by employing RDKit using ETKDG method and MMFF94 force field. Moreover, the 3D structures of chemicals were stored in various formats such as SDF, MOL, MOL2, PDB and PDBQT using Open Babel. Of the 3420 chemicals, we were able to obtain the 3D structure for 3417 chemicals. Moreover, the chemicals were mapped to identifiers in several databases including PubChem, ChEMBL^57,58^ (https://www.ebi.ac.uk/chembl/), ChEBI (https://www.ebi.ac.uk/chebi/), FDA SRS (https://precision.fda.gov/uniisearch), ChemSpider, Molport (https://www.molport.com/), SureChEMBL (https://www.surechembl.org/) and ZINC ^59–61^ (https://zinc15.docking.org/). Further, the chemicals were associated to published spectrum information obtained via different experimental techniques, by using compiled information in SpectraBase (https://spectrabase.com/) database.

For each chemical, we computed the natural product likeness score or NP-likeness^62^ score (https://github.com/rdkit/rdkit/tree/master/Contrib/NP_Score) using RDKit. Moreover, ClassyFire^63^ was used to obtain a hierarchical classification for the chemicals namely, kingdom, superclass, class, and subclass. Of the 3420 chemicals in our compilation, ClassyFire could not compute the classification for 49 chemicals. Further, physicochemical properties and drug-likeness properties of chemicals were computed using RDKit (https://www.rdkit.org/). Moreover, Absorption, Distribution, Metabolism, and Excretion (ADME) properties of chemicals were predicted using SwissADME.^64^ Of the 3420 chemicals in our compilation, SwissADME failed to compute the ADME properties for 13 chemicals. Furthermore, we computed 1875 molecular descriptors (both 2D and 3D descriptors) for each chemical using PaDEL software.^65^ Moreover, we computed the molecular scaffolds for the 3420 chemicals at graph/node/bond (G/N/B) level, graph/node (G/N) level, and graph level, using RDKit.^66,67^ G/N/B has connectivity, element and bond information, G/N has connectivity and element information, and graph level has only connectivity information.

### Chemical similarity network

The structural similarity between any two essential oil chemicals was ascertained by computing the pairwise Tanimoto coefficient (Tc)^68^ using the Extended Circular Fingerprints (ECFP4)^69^ in RDKit. Thereafter, a chemical similarity network was generated wherein the nodes are chemicals and two nodes are connected with an edge if they have Tc ≥ 0.5.

### Web interface and database management

We have encapsulated the compiled data in an online database namely, *DataBase of Essential oil Chemical profiles of Medicinal plants of India* (sCentInDB) with a user-friendly web interface to widely disseminate the compiled dataset on essential oil profiles of Indian medicinal plants and associated metadata (Figure 2). sCentInDB can be accessed at: https://cb.imsc.res.in/scentindb. In the back-end, we employed the open-source relational database management system MariaDB (https://mariadb.org/) to store the compiled data. Moreover, the front-end of the website was created using the open-source CSS framework Bootstrap 5.3.0 (https://getbootstrap.com/docs/5.3/) customized with in-house HTML, PHP (https://www.php.net/), JavaScript and jQuery (https://jquery.com/) scripts. Furthermore, JSME Molecule Editor^70^ has been incorporated into the web interface to enable drawing of chemical structures. sCentInDB is hosted on an Apache (https://httpd.apache.org/) webserver running on Debian 9.4 Linux Operating System.

**Figure 2:**
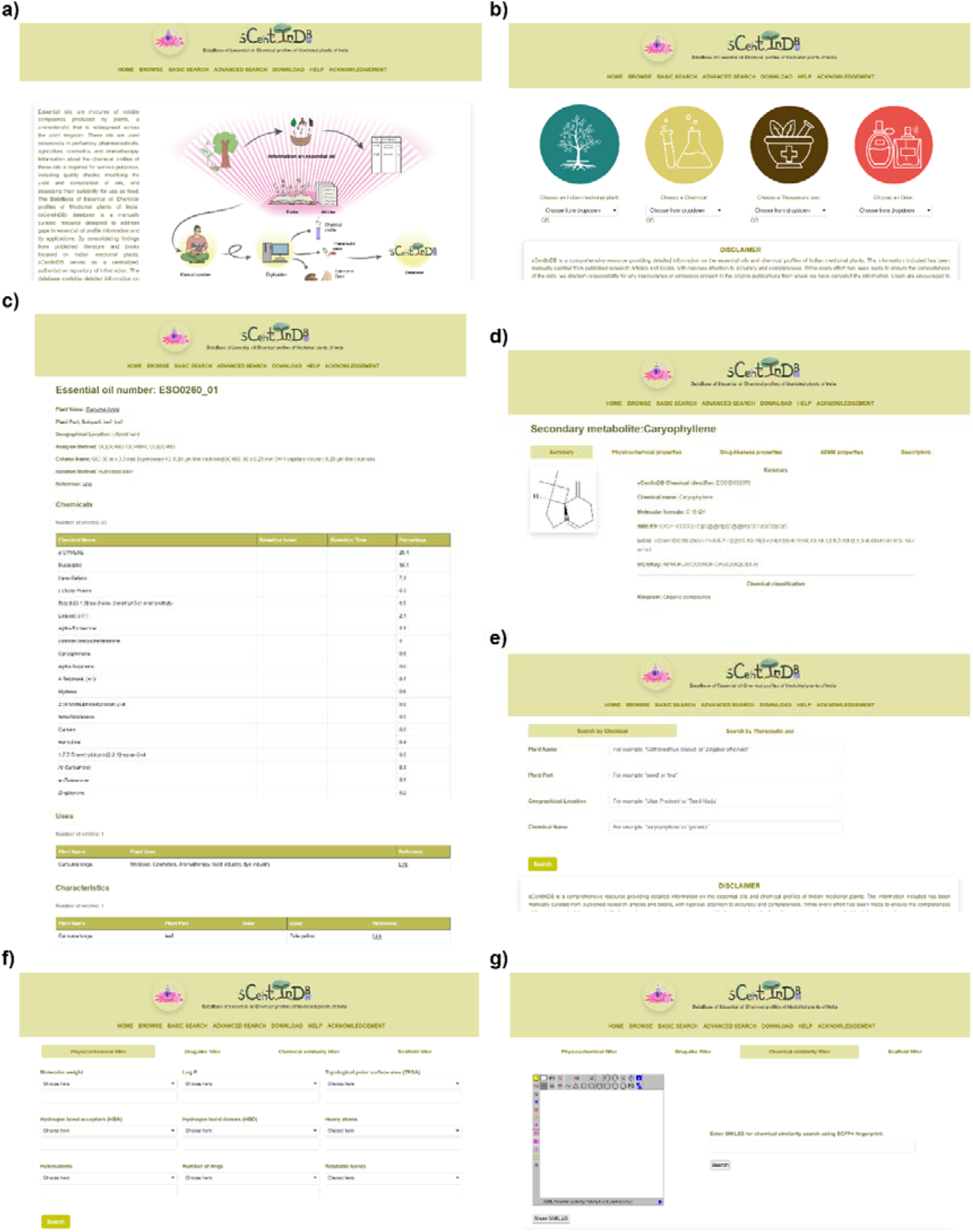
Snapshots of the web interface of sCentInDB. **(a)** Home page of sCentInDB providing a brief description of the database. **(b)** Browse page of sCentInDB showing four different ways for browsing the compiled information. **(c)** Snapshot of the detailed page on an essential oil. The displayed page corresponds to the leaf oil of the plant *Curcuma longa*, and contains information on its identifier, plant name, part name, analysis method, chemical composition, general uses, odor, and color. **(d)** Snapshot of the detailed page corresponding to a chemical with five different tabs for summary, physicochemical properties, drug-likeness properties, ADME properties, and descriptors. **(e)** Basic search page of sCentInDB displaying two different options to search the database. **(f)** Advanced search page of sCentInDB which enables filtering and retrieval of chemicals via four different options. **(g)** Chemical similarity filter option in the Advanced search page allowing users to retrieve structurally similar chemicals in the database based on a specific user query.

## Results and Discussion

### sCentInDB: A FAIR compliant DataBase of Essential oil Chemical profiles of Medicinal plants of India

We have built a user-friendly database namely, sCentInDB, which stands for DataBase of Essential oil Chemical profiles of Medicinal plants of India. The database contains compiled information on essential oils of Indian medicinal plants and is openly accessible for academic research at: https://cb.imsc.res.in/scentindb/. Users can access the compiled information on the website through three options namely, Browse, Basic search and Advanced search (Figure 2a). Figure S1 displays the Entity-Relationship Diagram (ERD) for the different entities in sCentInDB.

#### Browse

The Browse tab allows users to browse essential oil information in four different ways: Indian medicinal plant, chemical, therapeutic use, and odor (Figure 2b). By choosing an Indian medicinal plant from the drop-down menu via the ‘Choose an Indian medicinal plant’ option, the user will be redirected to a page with information on plant part, essential oil identifier, chemicals found in the oil, and the published literature reference. By clicking on the plant name, the user can open a detailed page about the plant. Similarly, by clicking on the oil identifier, the user will be redirected to a detailed page about the oil, including all the associated metadata for that oil (Figure 2c). By choosing a chemical from the drop-down menu via the ‘Choose a Chemical’ option, the user will be displayed a table containing the chemical identifier, essential oil profiles containing the chemical, and a link that redirects to a detailed page about the chemical. This includes summary information on the chemical, its physicochemical properties, its drug-likeness properties, its ADME properties, and its molecular descriptors (Figure 2d). By choosing a therapeutic use term from the drop-down menu via the ‘Choose a Therapeutic use’ option, the user will be displayed a page with information on essential oil at plant part level with the corresponding therapeutic use, therapeutic use identifiers in external databases, and link to published literature reference. By choosing an odor term from the drop-down menu via the ‘Choose an Odor’ option, the user will be displayed a page containing information on essential oil at plant part level with corresponding odor, uniformized odor terms and the link to published literature reference.

#### Basic Search

The basic search tab allows text-based search to retrieve the curated information on essential oil via two options namely, by chemical and by therapeutic use (Figure 2e). The ‘Search by Chemical’ option allows users to enter complete or partial name of plant, plant part, geographical location, chemical name, and chemical composition in percentage. This generates a table containing information on essential oil relevant to the user query. The ‘Search by Therapeutic use’ option enables users to search for therapeutic uses of essential oils at plant part level with link to published literature reference.

#### Advanced Search

The advanced search tab enables users to filter and retrieve a subset of chemicals in sCentInDB based on various features including physicochemical properties, drug-likeness properties, chemical similarity, and molecular scaffolds (Figure 2f). The ‘Physicochemical filter’ allows users to filter chemicals based on molecular weight, Log P, topological polar surface area (TPSA), number of hydrogen bond acceptors, number of hydrogen bond donors, number of heavy atoms, number of heteroatoms, number of rings, and number of rotatable bonds. The ‘Drug-like filter’ helps in retrieving chemicals based on a combination of various drug-likeness scoring schemes. The ‘Chemical similarity filter’ allows users to draw a chemical’s structure using the molecular editor or provide input in SMILES format, and thereafter, retrieve structurally similar chemicals present in sCentInDB (Figure 2g). The ‘Scaffold filter’ helps in retrieving chemicals based on molecular scaffold at Graph/Node/Bond (G/N/B), Graph/Node (G/N), or Graph levels.

#### Compliance with FAIR principles

While creating the database, we ensured that sCentInDB adheres to FAIR^71^ principles, which stands for Findable (F), Accessible (A), Interoperable (I), and Reusable (R). sCentInDB is compliant with the FAIR principles due to the following reasons:

- sCentInDB has a stable and permanent web address (F, A).
- All entries in sCentInDB can be accessed by stable URLs (F).
- Plants, chemicals, and therapeutic use terms are linked to unique external identifiers in standard databases (F, I).
- Data within sCentInDB can be accessed through Browse, Basic Search, and Advanced Search tabs by providing user query (A).
- Users do not need to create an account to access sCentInDB (A).
- sCentInDB has Help page explaining how to use the data (A).
- Data within sCentInDB is available in human readable format (html) and can be downloaded in machine readable format (.csv), and therefore, can be reused for further research (A, R).
- License of sCentInDB database also allows users to freely use the associated data for academic research (R).
- The data curation workflow is explained in detail in the associated publication (R).

### Exploration of Indian medicinal plants with documented essential oil profiles in sCentInDB

We compiled detailed information on essential oils from 554 Indian medicinal plants, including their chemical composition and reported therapeutic uses from published literature (Methods). Additionally, the Indian medicinal plants were annotated with information on their taxonomic classification, their use in traditional systems of medicine, and categorization in the IUCN Red List of threatened species. We observed that the 554 plants with essential oil profiles in sCentInDB belong to 86 taxonomic families. Figure 3a shows the taxonomic families with at least 5 plants in the database. Lamiaceae, commonly known as mint family, has the largest representation (95 plants) in sCentInDB. As one of the largest flowering plant families in the world, Lamiaceae holds significant economic importance in cosmetics, food, and medicine.^72,73^ Further, Asteraceae family has second largest representation (76 plants) in sCentInDB (Figure 3a). Moreover, Angiosperms or flowering plants constitute 96.2%, Gymnosperms account for 3.6%, and Pteridophytes comprise only 0.2% of the plants in sCentInDB (Figure 3b). Many plants in sCentInDB are documented to be used in traditional systems of medicine including 212 plants in Ayurveda, 187 plants in Siddha, 145 plants in Unani, 78 plants in Sowa Rigpa, and 60 plants in Homeopathy (Figure 3c). According to the IUCN Red list of threatened species, 6 plants in sCentInDB are classified as critically endangered and 13 are listed as endangered (Figure 3d).

**Figure 3:**
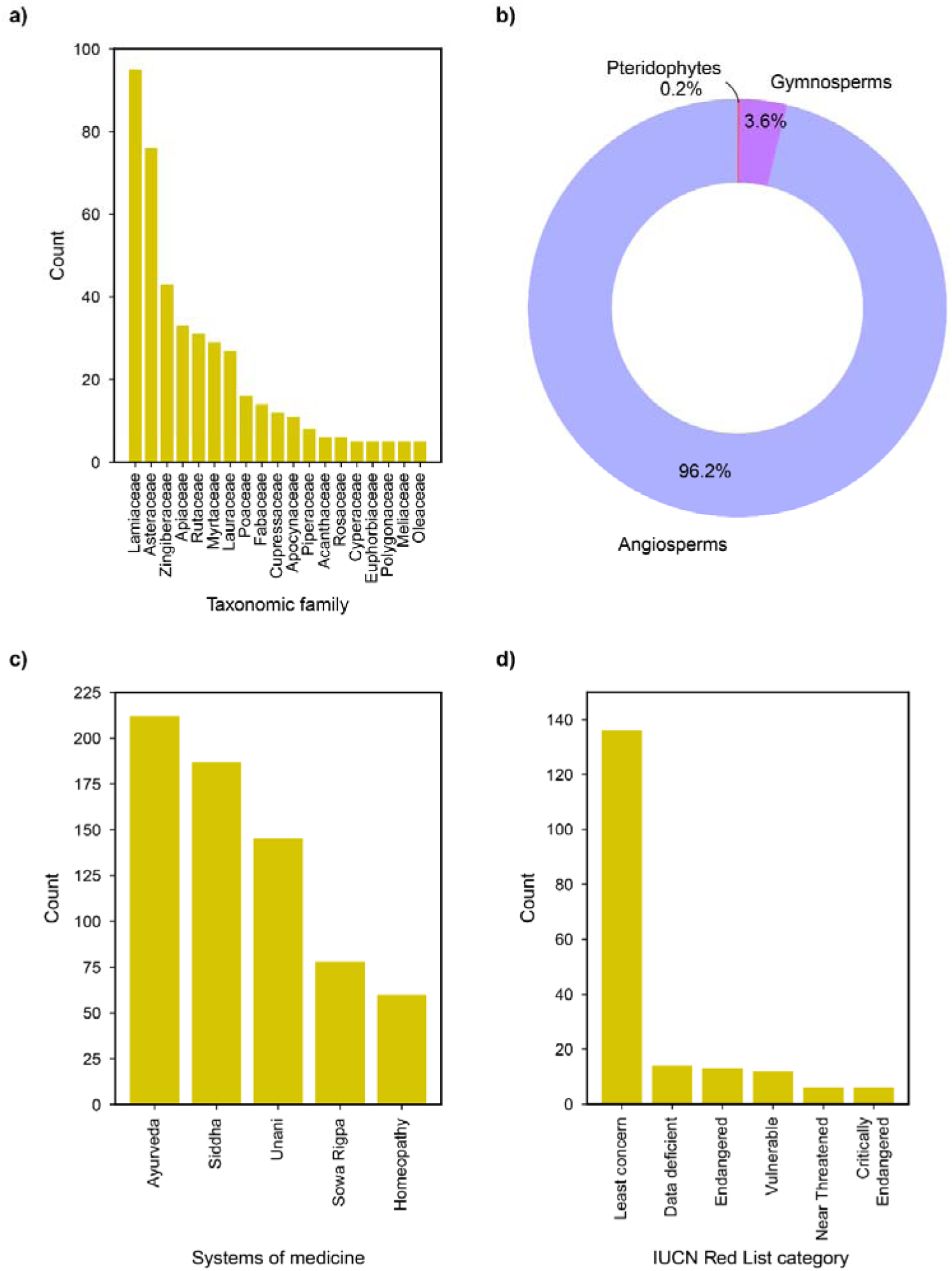
Basic statistics of medicinal plants in sCentInDB. **(a)** Bar plot depicting the number of plants across taxonomic families with at least five plant species in the database. **(b)** Pie chart showing the distribution of plants across groups in the database. **(c)** Bar plot depicting the distribution of plants across various traditional systems of Indian medicine. **(d)** Bar plot showing the distribution of plants across different IUCN Red List categories.

### Analysis of the essential oil information compiled in sCentInDB

sCentInDB compiles detailed information on essential oils of Indian medicinal plants, including their chemical composition, therapeutic uses, odor, and color, from published literature. Each essential oil was assigned a unique identifier, corresponding to the specific plant and the plant part from which it was derived. In addition, we assigned sample identifiers to each essential oil profile, taking into account variations in location, sampling conditions, and different published references, to ensure that different samples of the same essential oil across published literature were clearly distinguished. For example, in sCentInDB, the essential oil derived from the rhizome of the plant *Zingiber officinale* is assigned the identifier ‘ESO0822’. The profile for this oil is based on 88 samples which are denoted by sample identifiers ‘ESO0822_01’ to ‘ESO0822_88’, and capture the variations in the geographical locations from where the plant samples were sourced, sampling conditions and literature source. Altogether, sCentInDB catalogs 815 essential oils at the plant part level, which expands to 2170 when considering the multiplicity of samples reported in published literature. Among the different plant parts, leaf is the most used part for essential oil extraction, being utilized in a maximum number (266) of plants of the database (Figure 4a). Further, among the different Indian medicinal plants with essential oil profiles, the database compiles essential oils for maximum of six different plant parts for *Mentha arvensis* (Figure 4b). Moreover, the chemical caryophyllene is documented to be present in maximum number of essential oils (627) compiled in the database (Figure 4c).

**Figure 4:**
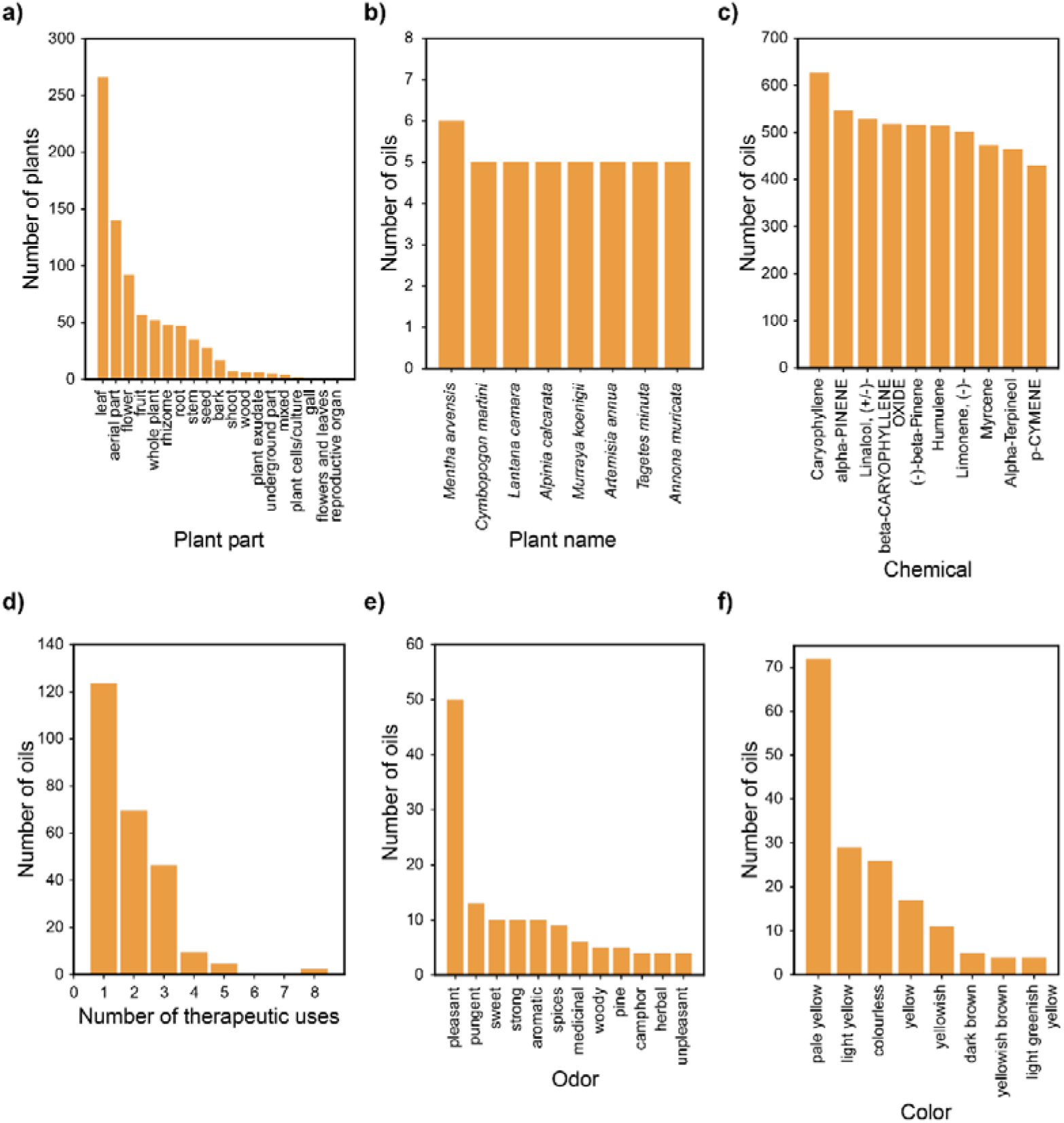
Basic statistics of essential oil data compiled in sCentInDB. **(a)** Bar plot showing the distribution of the plants across different plant parts in the database, from which the essential oils are derived. **(b)** Bar plot depicting the top Indian medicinal plants with documented information based on the number of essential oils at the level of plant parts. **(c)** Bar plot showing the top most represented chemicals in the essential oil data. **(d)** Bar plot depicting the distribution of the essential oils in terms of their number of therapeutic uses. **(e)** Bar plot showing the distribution of the essential oils across the most represented odor terms associated with the essential oils. **(f)** Bar plot showing the distribution of the essential oils across the most represented color terms associated with the essential oils.

#### Therapeutic uses of essential oils in sCentInDB

The therapeutic use information for the essential oils at the level of plant parts were compiled from published literature. Among the 554 essential oil-bearing plants in sCentInDB, we were able to compile the therapeutic use information for oils corresponding to 209 plants. In terms of essential oil, of the 815 oils in the database, we were able to collect the therapeutic use information for 253 oils. In order to standardize these therapeutic use terms, we mapped them to corresponding terms in Medical Subject Headings (MeSH) (https://www.ncbi.nlm.nih.gov/mesh/), Unified Medical Language System (UMLS) (https://www.nlm.nih.gov/research/umls/index.html), and International Classification of Diseases 11th Revision (ICD-11) (https://icd.who.int/browse/2024-01/mms/en). This resulted in a non-redundant list of 41 standardized therapeutic use terms and 471 plant-part-therapeutic use associations in sCentInDB database. We observed that majority of the oil in sCentInDB have only one therapeutic use term (Figure 4d), and anti-bacterial and antifungal activity are the most represented therapeutic use terms across the oils in sCentInDB.

#### Odor and color of essential oils in sCentInDB

The odor data of the essential oils at the level of plant parts was compiled from published research articles. Among the 815 oils in sCentInDB, we were able to compile odor data for 115 oils. To standardize these odor data, we primarily relied on OlfactionBase^74^, refining them to 54 uniformized odor terms. Notably, the odor of essential oils is often described using a combination of odor terms including the odor families, intensity etc. It is important to note that given an essential oil, different research papers may have reported its odor differently. This is because odor perception is a subjective psychophysical process.^75^ Additionally, variations in odor descriptions may arise due to differences in sampling conditions across published literature.^76^ We observed that the odor term ‘pleasant’ is associated with maximum number of oils (50) in sCentInDB (Figure 4e).

Similarly, the color data of the essential oils at the level of plant parts was compiled from published research articles. Among the 815 oils in sCentInDB, we were able to compile the color data for 210 oils. Thereafter, we manually uniformized the color data into 48 distinct color terms. Similar to the case of odor, the color of essential oil can have variations across published literature due to the differences in sampling conditions.^77,78^ We also observed that color term ‘pale yellow’ is associated with maximum number of oils (72) in sCentInDB (Figure 4f).

Altogether, sCentInDB has compiled chemical composition information for 2170 essential oils (including multiple samples for a particular oil) at plant part level. Additionally, it captures information on therapeutic use, odor and color information for essential oils from published literature.

### Analysis of the chemical space of essential oils documented in sCentInDB

sCentInDB compiles an *in silico*, non-redundant and stereo-aware library of 3420 unique chemicals found in the essential oils of Indian medicinal plants at the plant part level. To curate this chemical library, we obtained the 2D structures of these chemicals from multiple sources (Methods). For completeness, we also accounted for all chemicals contributing to the essential oil profile, even if their structures could not be retrieved. Furthermore, these chemicals were checked for structural similarity, and thereafter, unique structures were resolved based on their stereochemistry using InChI representation. This led to a non-redundant list of 3420 chemicals with their structural information, across 2170 essential oils (including oil samples) at the level of plant parts which accounts for 39438 plant-part-chemical associations. Figure S2a shows the distribution of the number of chemicals across Indian medicinal plants documented in sCentInDB. From the figure, it can be observed that there is one plant with 352 chemicals, *Jasminum sambac*, and 3 other plants produce 332 chemicals each. Interestingly, the top 20 plants in our database, in terms of the number of documented phytochemicals found in their essential oils, cover nearly one-third of the total phytochemicals in the database.

The phytochemicals in sCentInDB were given unique chemical identifiers, with each identifier corresponding to a chemical name, respective structural features and hyperlinks to external standard chemical databases. The 2D and 3D structures of the chemicals are provided in five different file formats (Methods). Further we computed the molecular scaffold(s), the core structure of a molecule, which has extensive use in the medicinal chemistry field. We have adapted the scaffold definition given by Lipkus *et al.*^66,67^ and computed the scaffold of the chemicals at three different levels (Methods).

#Figures S2b-g show the distributions of six physicochemical properties for the 3420 chemicals in sCentInDB. The chemicals in sCentInDB were classified into 18 superclasses, 103 classes, and 158 subclasses using ClassyFire (Figure 5a). Among these 18 superclasses, the top three superclasses are lipids and lipid-like molecules, organic oxygen compounds, and benzenoids comprising 1708, 455 and 371 chemicals, respectively in sCentInDB (Figure 5a). Furthermore, in order to obtain a natural product specific classification, we used NP classifier^79^ to classify the chemicals in sCentInDB based on their biosynthetic pathways. The top three biosynthetic pathways were terpenoids, fatty acids, and shikimates and phenylpropanoids corresponding to 1662, 1099 and 264 chemicals, respectively in sCentInDB (Figure 5b).

**Figure 5:**
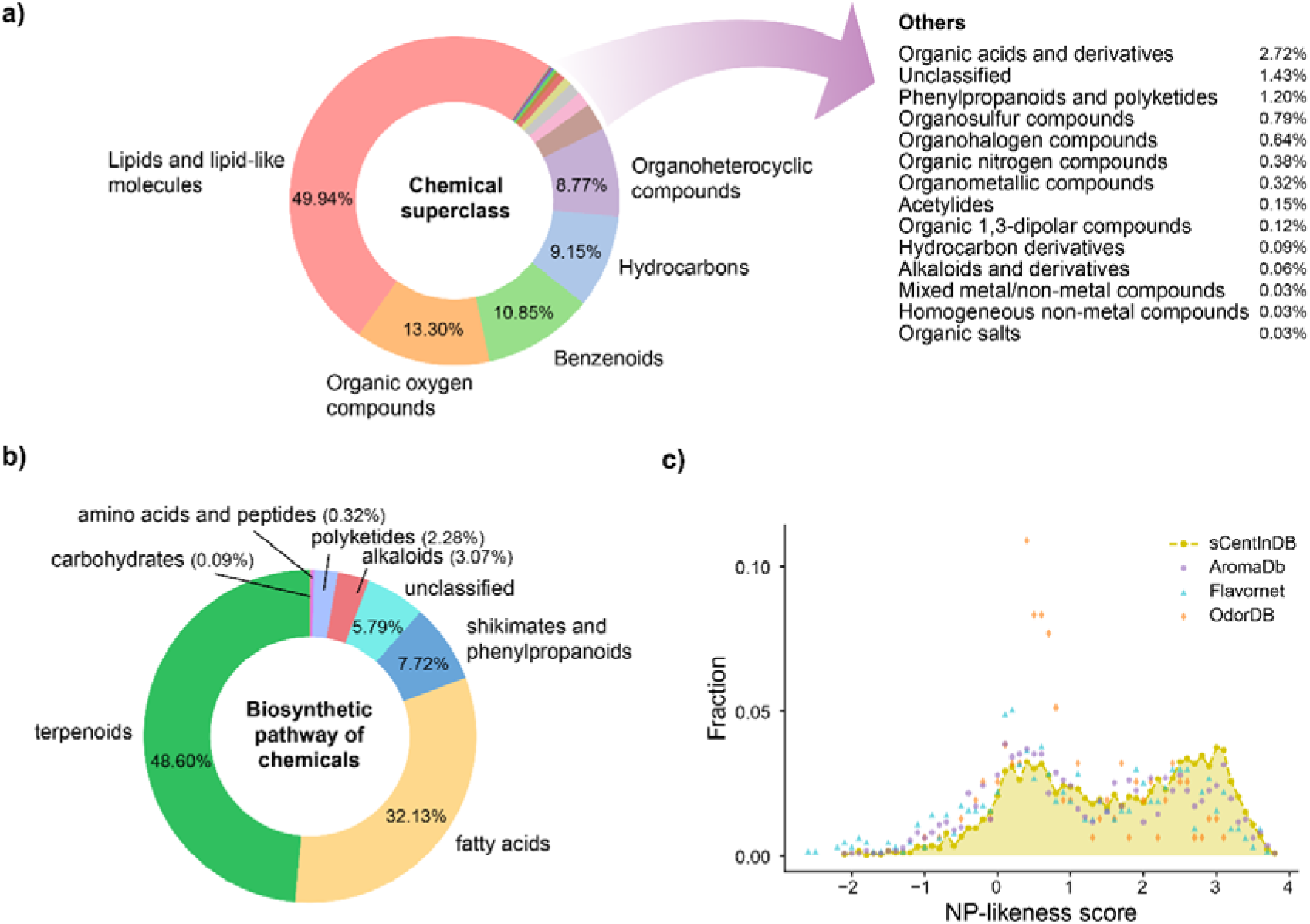
Chemical superclass, biosynthetic pathways, and natural product-likeness scores for 3420 chemicals in sCentInDB. **(a)** Distribution of the chemical superclass for chemicals as predicted by ClassyFire. **(b)** Distribution of the biosynthetic pathways of chemicals as predicted by NP classifier. **(c)** Distribution of NP-likeness scores of chemicals in sCentInDB as compared to other odor or aroma molecule libraries.

NP-likeness^62^ score is a measure to quantify the natural product likeness of a chemical. This score ranges from -5 to +5, wherein the higher the score for a chemical, the more likely for it to be a natural product.^62^ In general, NP-likeness scores of natural product libraries are predominantly positive.^80,81^ As expected, >90% of the chemicals in sCentInDB have NP-likeness scores to be positive, as the database is specific for natural products. Figure 5c shows the distribution of NP-likeness scores for chemicals in sCentInDB. It can be seen that this distribution is skewed toward the positive side, and similar to the distribution for other chemical libraries containing aroma molecules (Figure 5c).

#### Chemical similarity network

We generated the chemical similarity network (CSN) of the 3420 chemicals in sCentInDB to analyze their structural diversity (Methods; Figure 6). We observe that the CSN has 183 connected components (with two or more chemicals each) and 618 isolated nodes. Notably, the largest connected component of the CSN consists of 1452 chemicals, with the majority of these chemicals (947 of 1452) produced through fatty acid biosynthetic pathway. Interestingly, among the 1099 chemicals belonging to the fatty acid biosynthetic pathway in the database, 947 are clustered into a single component in the CSN, while chemicals belonging to the terpenoid biosynthetic pathway are more dispersed across the CSN, indicating greater structural diversity of terpenoids^82,83^. Additionally, within the largest connected component of the CSN, *Jasminum sambac* is documented to produce the highest number of chemicals (285), the majority of which belong to the fatty acid biosynthetic pathway.

**Figure 6:**
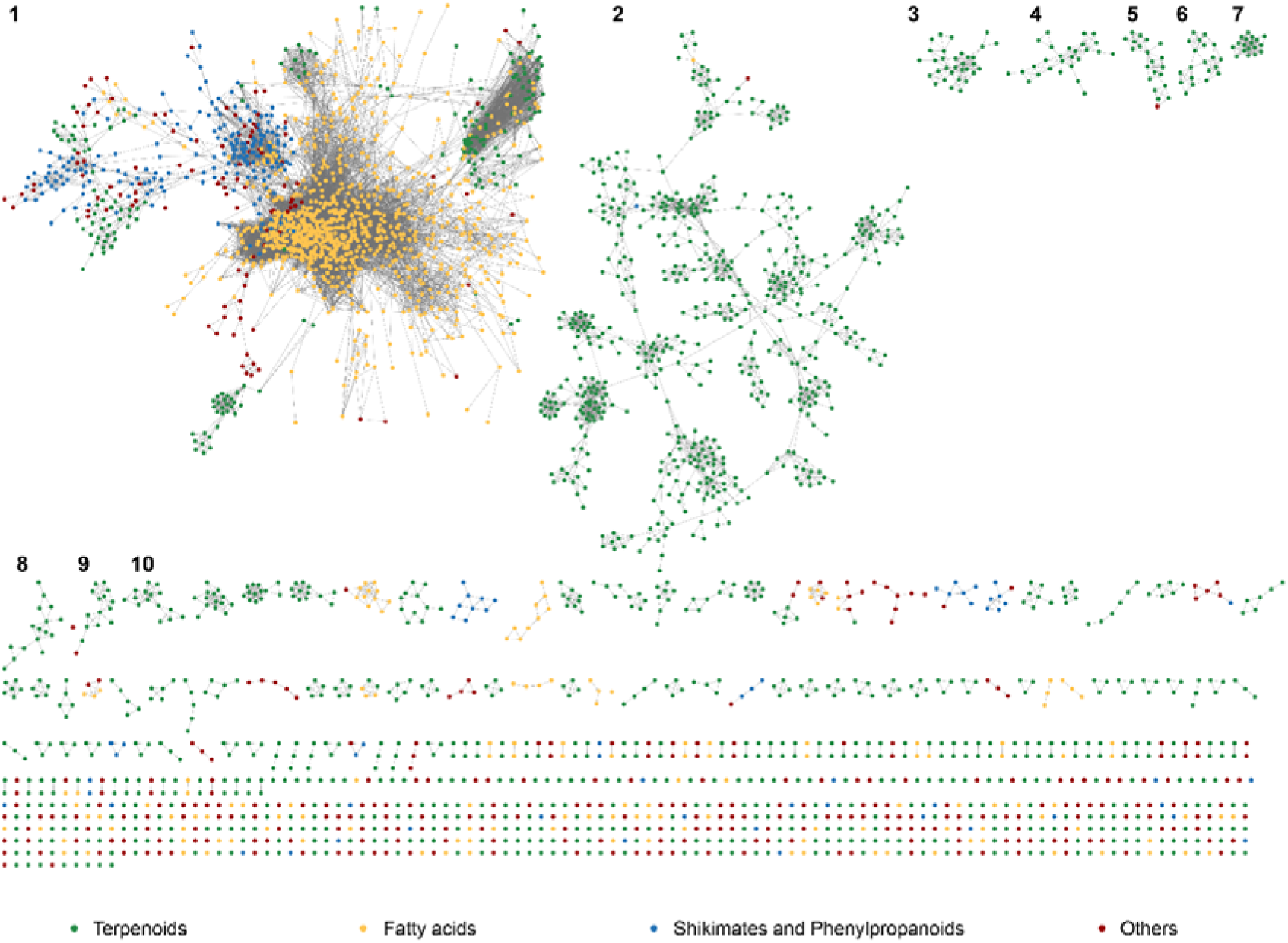
Chemical similarity network (CSN) comprising of the 3420 chemicals in sCentInDB. This network visualization was generated using Cytoscape.^96^ The nodes in the network correspond to the chemicals, and two nodes are connected if the chemical similarity computed using the Tanimoto coefficient (Tc) is ≥ 0.5. The nodes in the CSN are colored based on their biosynthetic pathway predicted by NP classifier: green for terpenoids, yellow for fatty acids, blue for shikimates and phenypropanoids, and dark red for others. In this CSN visualization, the top 10 connected components are numbered.

#### Molecular scaffold based diversity analysis

Molecular scaffold based diversity analysis of a chemical space can help identify small molecules with novel scaffolds of interest.^84,85^ We followed Lipkus *et al.*^66,67^ to compute Bemis and Murcko scaffolds^86^ at three different levels namely, graph/node/bond (G/N/B) level, graph/node (G/N) level, and graph level (Methods). We find that the chemical space of sCentInDB comprise 693 scaffolds at G/N/B level, 479 scaffolds at G/N level, and 362 scaffolds at graph level. We compared the scaffold diversity of sCentInDB with three other chemical libraries containing aroma molecules namely, AromaDb, Flavornet, and OdorDB. At the G/N/B level, sCentInDB is second only to AromaDb in terms of the fraction of scaffolds per molecule (N/M) and the fraction of singleton scaffolds per molecule (N_sing_/M) (Table 1). sCentInDB and AromaDb have very similar AUC values, suggesting similar scaffold diversity for both the essential oil libraries. The higher AUC values for sCentInDB and AromaDb in comparison to other natural product libraries^54,84^ can be attributed to the fact that essential oil chemicals are produced by a specific set of pathways and from a limited number of precursors.^87^

**Table 1:** Comparison of the scaffold diversity of sCentInDB with other chemical libraries containing aroma molecules. Note the scaffolds for each chemical library are computed at G/N/B level.

| Chemical library | Number of molecules (M) | Number of scaffolds (N) | $N_{\text{sing}}$ | N/M | $N_{\text{sing}}/M$ | $N_{\text{sing}}/N$ | AUC | $P_{50}$ |
| --- | --- | --- | --- | --- | --- | --- | --- | --- |
| sCentInDB | 3414 | 695 | 471 | 0.204 | 0.138 | 0.678 | 0.876 | 0.576 |
| OdorDB | 156 | 20 | 12 | 0.128 | 0.077 | 0.6 | 0.891 | 5.263 |
| Flavornet | 633 | 87 | 51 | 0.137 | 0.081 | 0.586 | 0.899 | 2.326 |
| AromaDb | 1109 | 239 | 167 | 0.216 | 0.151 | 0.699 | 0.874 | 0.84 |

To further analyze the structural diversity of the four chemical libraries, we plotted the cyclic system retrieval (CSR) curves for the scaffolds computed at the three levels. We observed that the CSR curves follow similar trend across the four libraries considered here (Figure 7). However, the CSR curve for OdorDB shifts closer to the diagonal line compared to other libraries as one moves from G/N/B to graph level (Figure 7). In an ideal scenario, the CSR curve of a chemical library, wherein each chemical has a unique scaffold, will correspond to the diagonal line. Similarly, the CSR curve of a chemical library, wherein scaffolds are evenly distributed among the chemicals, will also correspond to the diagonal line. This suggests the need for multiple metrics to fully assess the scaffold diversity.^88^ Figure 8 is a molecular cloud visualization of the scaffolds that occur in five or more chemicals in sCentInDB. We observed that the benzene scaffold dominates sCentInDB and is present in 387 chemicals.

**Figure 7:**
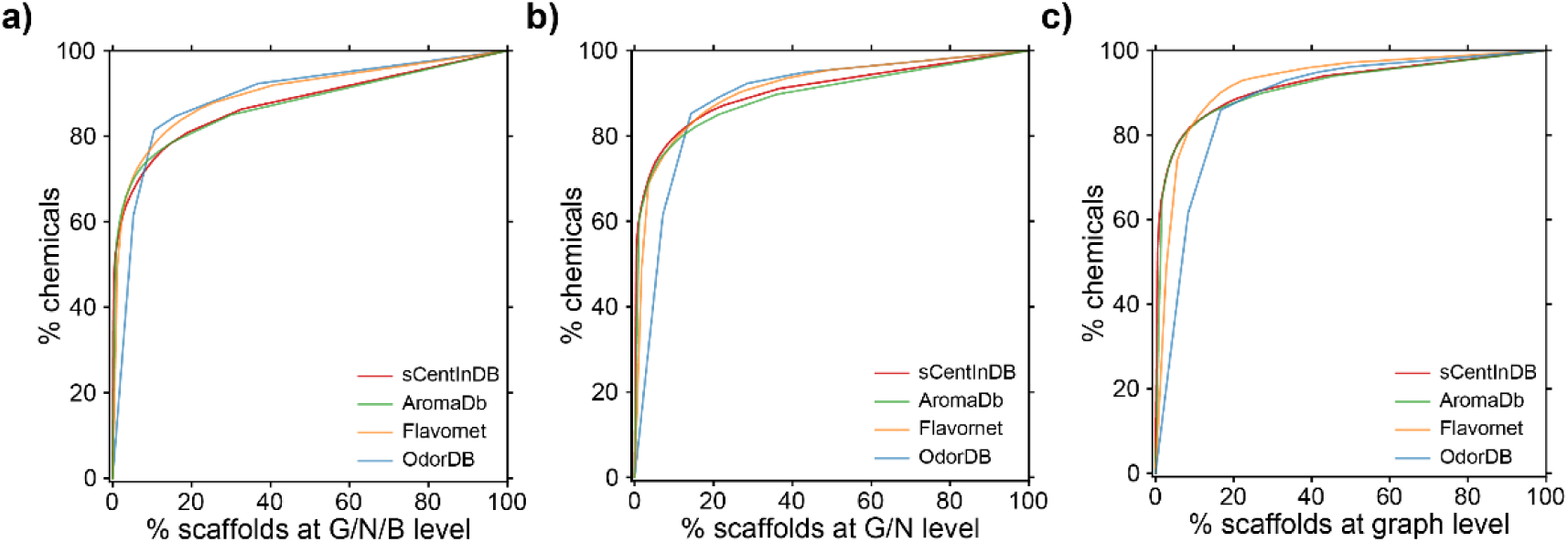
Cyclic system retrieval (CSR) curves comparing the scaffold diversity across four chemical libraries containing aroma molecules namely, sCentInDB, AromaDb, Flavornet, and OdorDB. The x axis of the CSR curve is the percent of scaffold and the y axis is the percent of compounds that contain those scaffolds. **(a)** CSR curves for scaffolds at G/N/B level. **(b)** CSR curves for scaffolds at G/N level. **(c)** CSR curves for scaffolds at graph level.

**Figure 8:**
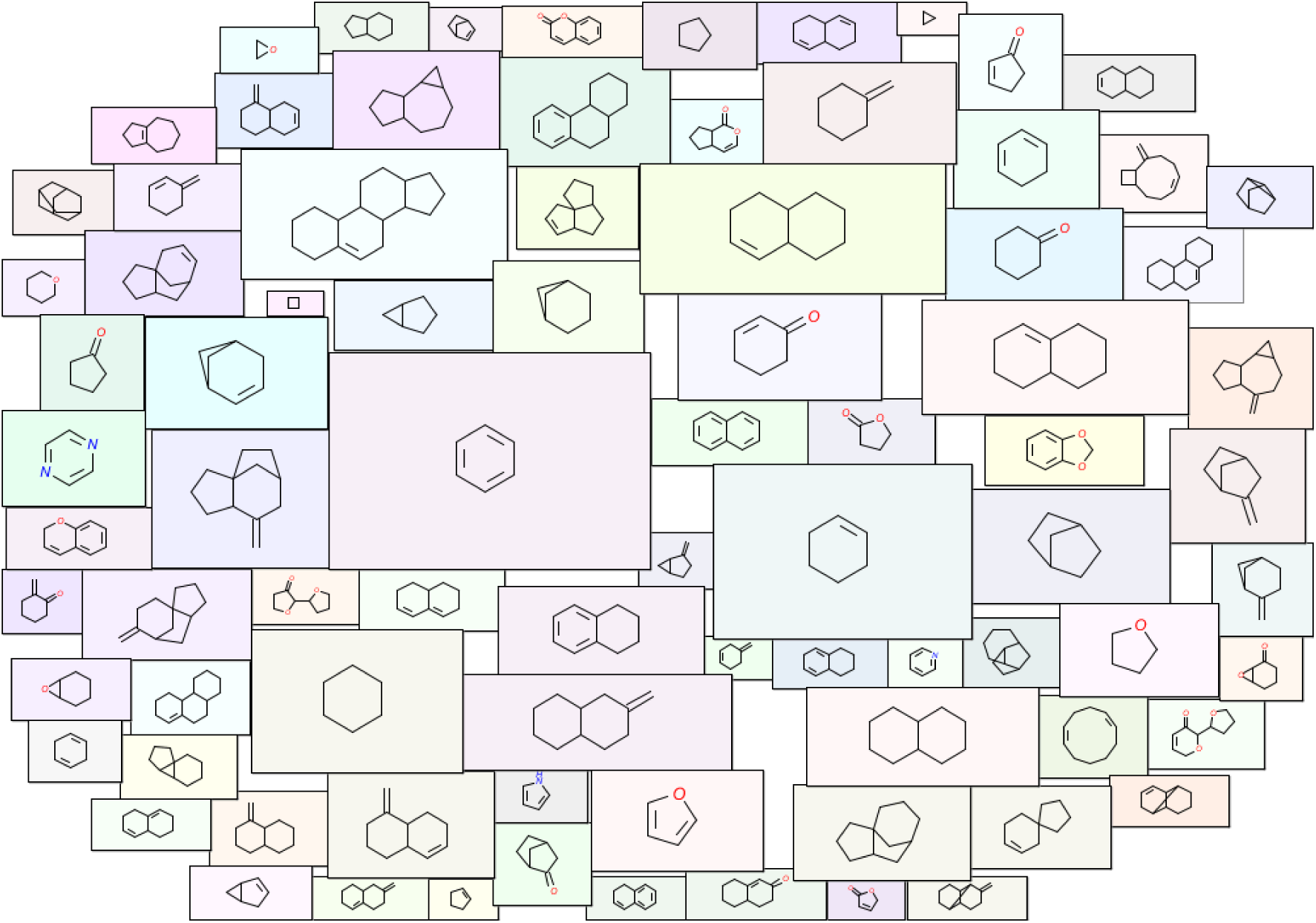
Molecular cloud visualization of top scaffolds at the G/N/B level which are present in at least five chemicals in sCentInDB. In this visualization, the size of each scaffold image is proportional to its frequency of occurrence in the 3420 chemicals in the database.

### Comparison of sCentInDB with related databases

We performed a comparative analysis of sCentInDB against previously built related databases. Based on this analysis, we find sCentInDB to be the most comprehensive resource for essential oils from Indian medicinal plants. Briefly, a few resources have been built previously to provide information on essential oils, and their chemical constituents. Among them, EssOilDB^4^ and AromaDb^52^ (https://aromadb.cimapbioinfo.in/) were developed to provide detailed information about essential oils, chemical composition, and associated properties. However, the website of EssOilDB is no longer functional, while AromaDb has limited coverage in terms of essential oils and medicinal plants in comparison to sCentInDB (Table 2). Moreover, other related databases including SuperScent^89^, OdorDB (https://ordb.biotech.ttu.edu/OdorDB/), and Flavornet (https://www.flavornet.org/) focus on flavors and odor molecules rather than on essential oils. Furthermore, proprietary resources such as EOUdb (https://essentialoils.org/plans) and AromaWeb (https://www.aromaweb.com/), capture information on essential oil profiles and aromatherapy, but are not freely accessible for academic research, with EOUdb requiring login credentials. In Table 2, we present a detailed comparison of three resources, sCentInDB, AromaDb and AromaWeb. From this comparative analysis, sCentInDB emerges as the most comprehensive resource to date for essential oil profiles of Indian medicinal plants and associated properties.

**Table 2:**
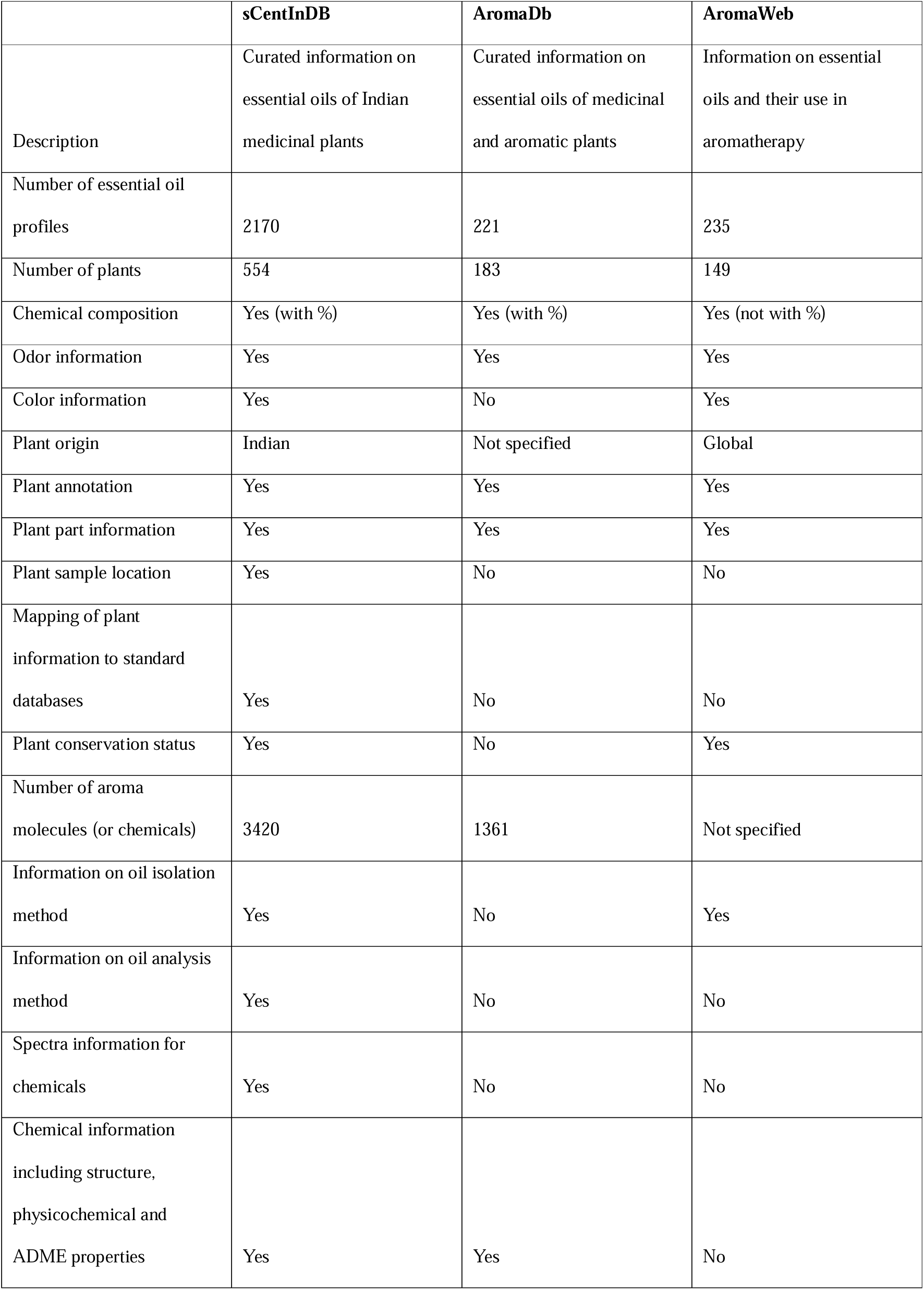
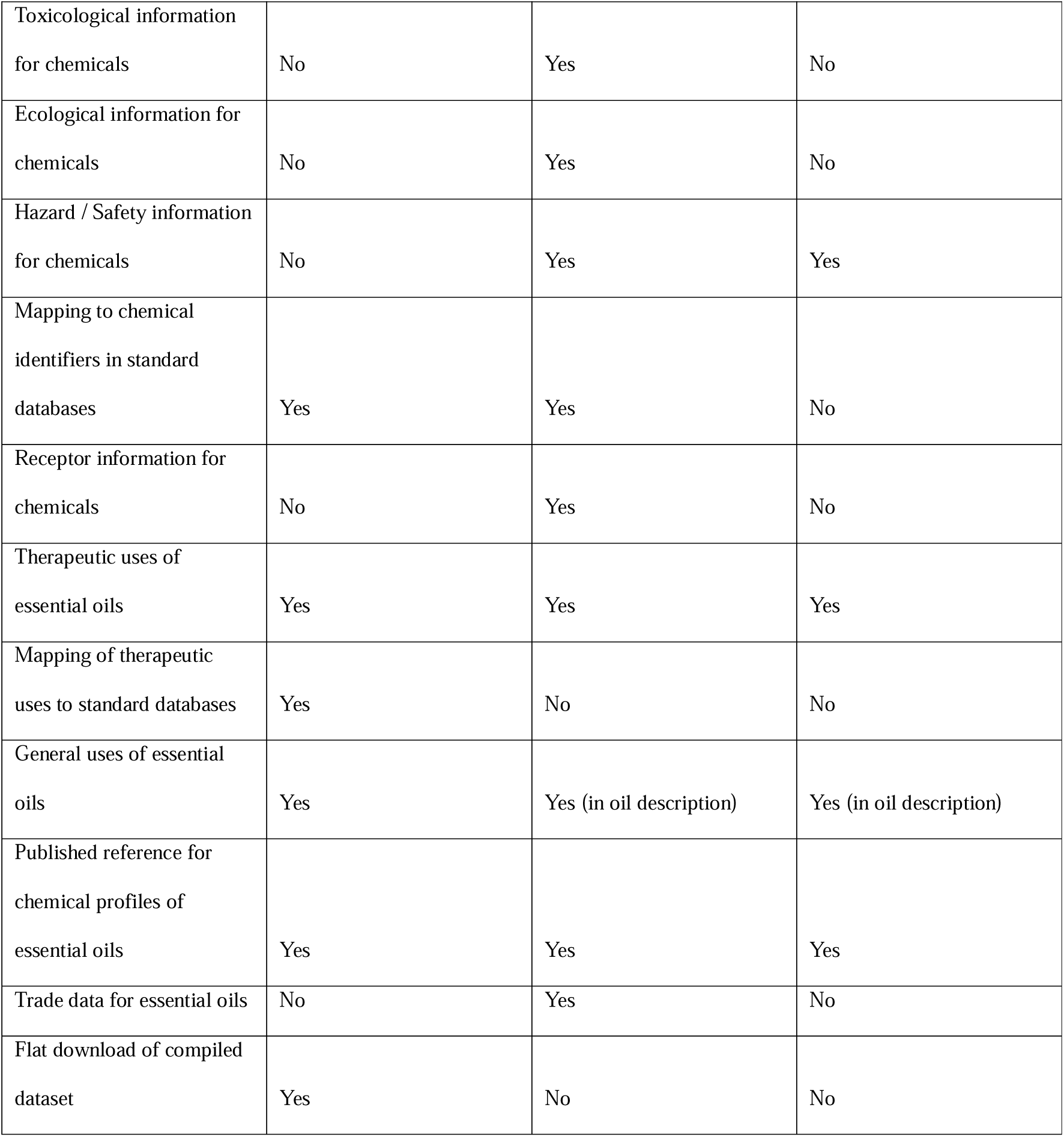
Comparison of the features of sCentInDB with related databases namely, AromaDb and AromaWeb.

## Conclusions

sCentInDB is an elaborate resource encapsulating the essential oil chemical profiles of Indian medicinal plants. It contains manually curated information on 2170 oil profiles at the plant part level, along with extensive metadata. Additionally, we have compiled data on the therapeutic uses, odor, and color of oils at the plant part level. To enhance interoperability, we have mapped and hyperlinked plant names, chemical names, and therapeutic use terms to other standard databases, thus complying with FAIR principles. Furthermore, we analyzed the structural diversity of the chemicals in sCentInDB. From the chemical similarity network (CSN), we observed that terpenoids exhibit greater diversity than chemicals from other biosynthetic pathways. Finally, we analyzed the scaffold diversity of the chemical space in sCentInDB using Cyclic System Retrieval (CSR) curves, which revealed that aroma molecule libraries derived from plants typically have less diversity due to the limited number of biosynthetic pathways from which they can be synthesized.

The compilation of essential oil compositions of aromatic plants is important for both research and industrial applications. In particular, this knowledge is required when genetically modifying plants to alter the yield and composition of the oil^40^. Further, many plants used for essential oil extraction are sourced from the wild, and therefore, stand the risk of extinction^90,91^. Genetic engineering offers a viable alternative by enhancing oil yield and modifying composition to achieve desired properties. In order to achieve these goals, a comprehensive understanding of the chemical composition of essential oils is necessary. Prior to our work, there were a few databases containing such information. However, previous databases are specialized libraries compiled from different perspectives, and thus, have limitations. Moreover, no existing databases on essential oils specifically focus on Indian medicinal plants. Thus, sCentInDB aims to fill this gap, serving as a valuable resource for researchers working with Indian medicinal plants in therapeutic, perfumery, and cosmetic industries.

Environmental factors^30,31^ and developmental stage^32,33^ of a plant have influence on the composition of essential oils. However, sCentInDB does not include this information. Such data could have been valuable in determining the optimal conditions and growth stages for cultivating plant parts for essential oil extraction. Additionally, the database does not contain information on recipes or combination of oils used in aromatherapy. While this was not the aim of the work, such data would be beneficial from a therapeutic perspective. Furthermore, trade-related information, such as the import and export data, was not collected, though it would have provided useful insights from a commercial standpoint.

Our database will be a highly valuable resource for the perfumery, pharmaceutical, fragrance, cosmetic, and food flavoring industries. It also enables users to find plants capable of producing their molecule of interest in high quantities. Although the structural diversity of essential oil chemicals is less in general^87^, unique chemical entities distinct from chemical libraries of other sources can be expected due to their unique properties. Essential oils have been researched for their application as complementary therapy^92,93^, and for reducing the side effects of modern medicine^94^. Additionally, the chemical composition of the plants can give insights into their genetic makeup^2^, making sCentInDB a valuable resource for systems biology research^95^. The data can be also used to predict the odor of new mixture of chemicals using machine learning models. Lastly, the region-specific data compiled in sCentInDB can help explore the relationship between regional climate conditions and their impact on oil composition.

## Author Contributions

**Shanmuga Priya Baskaran:** Conceptualization, Data Curation, Formal Analysis, Methodology, Software, Visualization, Writing – original draft, Writing – review & editing; **Geetha Ranganathan:** Conceptualization, Data Curation, Writing – original draft; **Ajaya Kumar Sahoo:** Conceptualization, Data Curation, Formal Analysis, Methodology, Visualization, Writing – original draft, Writing – review & editing; **Kishan Kumar:** Conceptualization, Data Curation; **Jayalakshmi Amaresan:** Data Curation, Writing – original draft; **Kundhanathan Ramesh:** Data Curation; **R.P. Vivek-Ananth:** Conceptualization, Data Curation; Writing – review & editing; **Areejit Samal:** Conceptualization, Formal Analysis, Methodology, Supervision, Writing – original draft, Writing – review & editing.

## Acknowledgements

The authors thank Gokul Balaji Dhanakoti for computational assistance and Gaurav Kumar for assisting in data curation. Areejit Samal would like to acknowledge funding from the Department of Atomic Energy (DAE), Government of India through the Apex Project to The Institute of Mathematical Sciences (IMSc), Chennai, India. The funders have no role in the study design, data collection, data analysis, manuscript preparation, or decision to publish.

