## Supplementary material for "sCentInDB: A database of essential oil chemical profiles of Indian medicinal plants": SI Figure

### Supplementary Figures S1-S2

### for

#### sCentInDB: A database of essential oil chemical profiles of Indian medicinal plants

Shanmuga Priya Baskaran[a,b], Geetha Ranganathan[a], Ajaya Kumar Sahoo[a,b], Kishan Kumar[a], Jayalakshmi Amaresan[a], Kundhanathan Ramesh[a], R.P. Vivek-Ananth[a,b], Areejit Samal[a,b,*]

[a] *The Institute of Mathematical Sciences (IMSc), Chennai 600113, India*

[b] *Homi Bhabha National Institute (HBNI), Mumbai 400094, India*

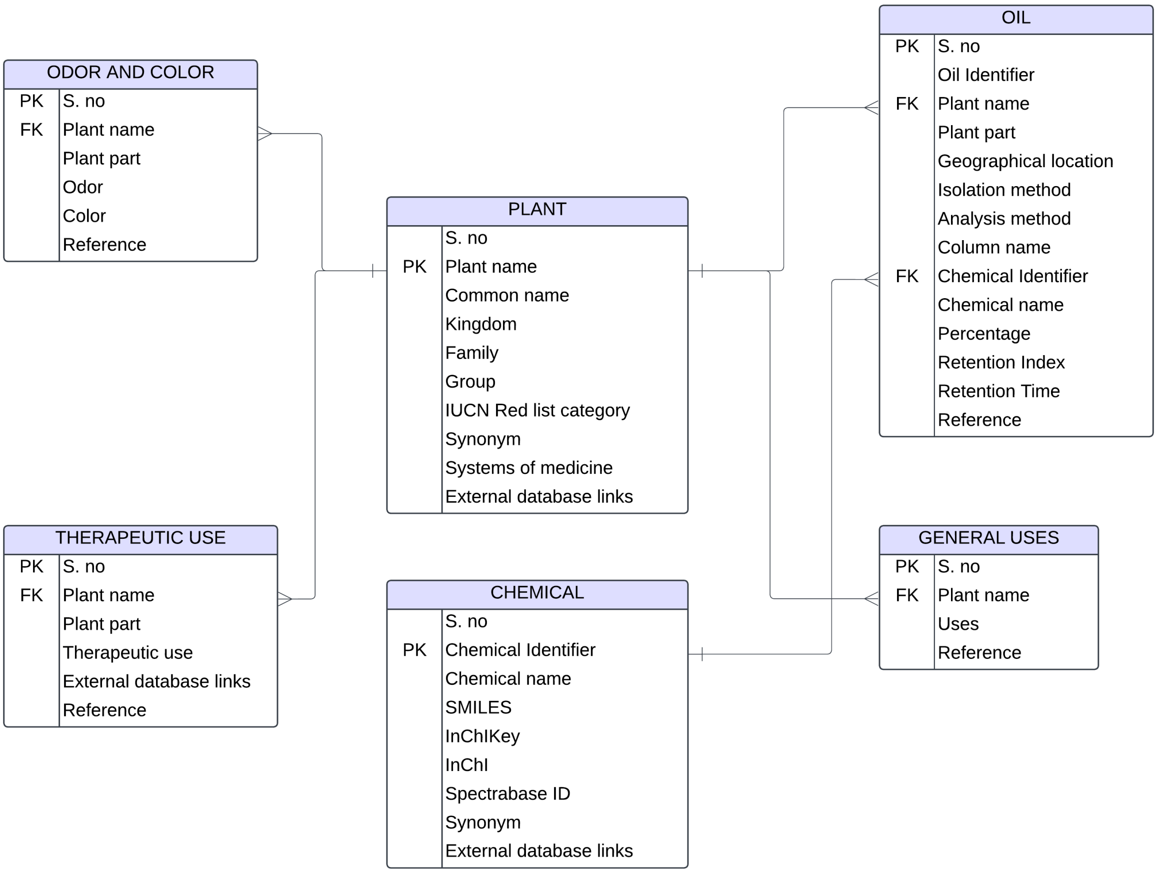


**Figure S1:** The Entity-Relationship Diagram (ERD) for sCentInDB database illustrating the relationships between various attributes across different entities. The Plant table serves as the primary entity, establishing connections with other entities such as Oil, Odor and Color, Therapeutic use, and General Uses. Additionally, the Chemical entity is connected with the Oil entity through the Chemical Identifier attribute.

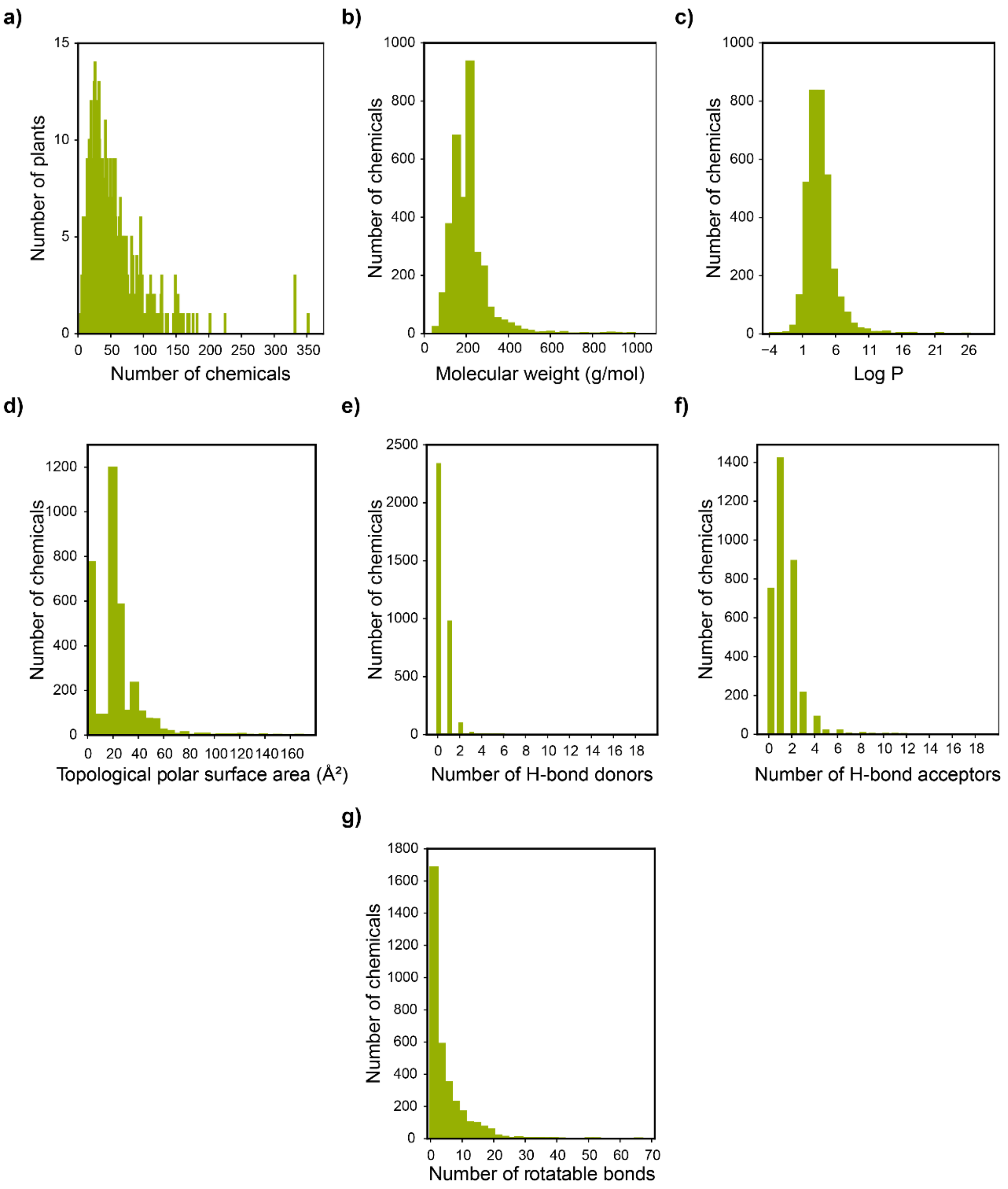


**Figure S2:** Basic statistics and physicochemical properties of essential oil chemicals in sCentInDB. **(a)** Distribution of Indian medicinal plants based on the number of associated chemicals. Distribution of six important physicochemical properties for the 3420 chemicals in sCentInDB: **(b)** molecular weight in g/mol, **(c)** Log P, **(d)** topological polar surface area in Å$^2$, **(e)** number of hydrogen bond donors, **(f)** number of hydrogen bond acceptors, and **(g)** number of rotatable bonds.
